# A structured evaluation and benchmarking of Boolean logical modelling tools

**DOI:** 10.1101/2025.10.28.684998

**Authors:** Benjamin Saalfeld, Emanuel Lange, Jacob Krüger, Petra Lutter, Robert Heyer

## Abstract

Biomolecules form complex interaction systems at different biological levels. The dynamics behind the interactions of biomolecules result in cellular phenotypes and responses. Two typical biological interaction systems are signal transduction and gene regulatory networks. Boolean models characterise these systems’ components with binary state variables and their interactions with rules governing their influence on each other, without requiring kinetic and abundance information about these components. Tools are available to define, analyse, and manage these models, but there is a lack of an overview and evaluation of such tools’ quality and functionality.

This work aims to facilitate the selection of software tools for Boolean modelling of signal transduction and gene regulatory networks by developing benchmark and evaluation criteria and applying them to the available tools. Twenty Boolean modelling tools were identified, which were assessed on their compliance to ten FAIR4RS (Findability, Accessibility, Interoperability, and Reusability for Research Software) aspects. Additionally, usability was assessed using a set of 25 developed sub-criteria. Furthermore, the functionalities were analysed, validated and the tools’ abilities to simulate and analyse Boolean models of different sizes were benchmarked.

The evaluation demonstrated that, whilst the results produced by the 20 tools were completely consistent, their analytical functionalities and capabilities to simulate larger models varied widely. Size constraints for tools’ functionalities ranged between 25 and more than 10,000 nodes. Each tool only covered part of the available functionalities, and many exhibited deficiencies in FAIR4RS compliance and usability. Based on our analysis, we see great potential for researchers to improve the usability of their tools along the FAIR4RS aspects in the future, with average tools in our evaluation not yet addressing 40% of the aspects (20% for the best-performing one). The aim is to present a useful overview of usability and functionality of Boolean modelling tools and to show how they can be improved throughout the entire community.

**Author summary:** Our study provides a set of criteria and a benchmarking toolkit for evaluating FAIR4RS and usability aspects as well as functionalities of Boolean modelling tools. We establish an overview of a wide range of Boolean modelling tools for signal transduction and gene regulatory network modelling, which is intended to make it easier for users to choose software according to their needs.

This work empowers developers by offering crucial assets: a standardised evaluation criteria set for comparing their tool’s functionalities, usability, and FAIR4RS compliance throughout the community. Our models and benchmarking assets further enable developers to precisely determine and improve their software’s model size limitations.

## Introduction

Systems biology aims to understand the functionality of biological systems on all levels by examining the properties of their components and interactions [1]. This interdisciplinary field employs a blend of experimental methodologies and computational tools to attain perspectives on the dynamics of biological systems [2]. Two pivotal biological systems regulating cellular processes ensuring appropriate responses to extracellular and intracellular signals are signal transduction and gene regulatory networks [3]. Understanding the dynamic behaviour of these networks is crucial for unravelling complex cellular behaviours, identifying potential drug targets for various diseases or finding missing interactions between signalling proteins [4].

Boolean models have emerged as a powerful concept for studying signal transduction networks, offering a comprehensive understanding of system dynamics and aiding the discovery of novel intervention strategies, particularly when detailed quantitative data is limited or unavailable [5]. Other modelling approaches, like ordinary differential equation models, require knowledge of the kinetic dynamics and abundances of the interacting components. This information is rarely accessible. Boolean models can be applied without these parameters [6]. While lacking the complexity of approaches requiring detailed knowledge, studies have shown the predictions of Boolean models to be useful due to the monotonous nature of the kinetics involved in gene regulation and signal transduction [7, 8]. By capturing essential regulatory interactions and dynamics, Boolean models enable researchers to explore various scenarios, predict cellular responses, and uncover novel regulatory mechanisms [2].

Today, many tools for Boolean modelling are available. However, the procedures for creating and analysing such models have not been standardised so far. There are three tools with related repositories for sharing Boolean models, the Cell Collective, BioModels, and GINsim [9–11]. Tools can differ in their analytical functionalities and in other aspects that influence usability, such as software accessibility and maintenance, or comprehensive documentation. Current implementations use varying data formats and definitions for models (Boolsim, GINML, BoolNet, SBML-qual, Truthtable…), potentially making them incompatible with each other, although efforts for standardisation have begun [12]. Evaluations of tools’ compatibility and usability help users choose software. To the best of our knowledge, there is currently no overview of Boolean modelling tools or a definition of standards for Boolean modelling.

To assess the differences between tools and compare them analytically, specific criteria must be provided. While some endeavours have been undertaken for standards like FAIR4RS (FAIR for Research Software), their criteria exclude many aspects of user comfort and focus more on software metadata and machine readability [13]. Concrete criteria must be employed to comprehensively and precisely review the different metrics in which Boolean modelling tools differ.

However, no study has yet provided a systematic overview that evaluates and compares these tools based on standardised criteria for functionality and usability. Therefore, a comprehensive assessment would greatly benefit users and developers in this field. In this study, we adopt a multifaceted approach, combining qualitative evaluation of tool features with quantitative performance benchmarking. Specifically, we evaluate the tools against the FAIR4RS principles and a self-developed usability assessment framework. This framework drew initial inspiration from Scott Jr. et al. 2023 “A structured evaluation of genome-scale constraint-based modelling tools for microbial consortia” but we developed a distinct set of criteria and introduced sub-criteria to better suit our evaluation goals and improve measurability [14]. For the quantitative assessment, we create and apply a benchmark to systematically test the performance and scalability of core analytical functionalities across a range of small-, medium-, and large-scale Boolean models. Ultimately, based on these qualitative and quantitative findings, we will provide recommendations for the most suitable modelling tools for specific applications, as well as suggestions for future development and improvement of the software.

## Results

We constructed an evaluation workflow including a usability evaluation, a FAIR evaluation, a functional evaluation, and a computational performance benchmarking (Fig 1). Resources and scripts to recreate the workflow can be found in the supporting information (S1 Code, S2 Code, S1 Data and S1 Text). As a first step, a search for tools was conducted yielding 20 tools (S1 Table). Next, minimum usage requirements as well as the usability criteria were developed, which were later used for evaluation of the tools (Fig 2). The tools were then further analysed based on FAIR4RS criteria. In the next step the functionalities of the tools were evaluated. Finally, the size constraints for the analysable models within the tools were benchmarked, and the calculation time, depending on the model size, was measured.

**Fig 1.**
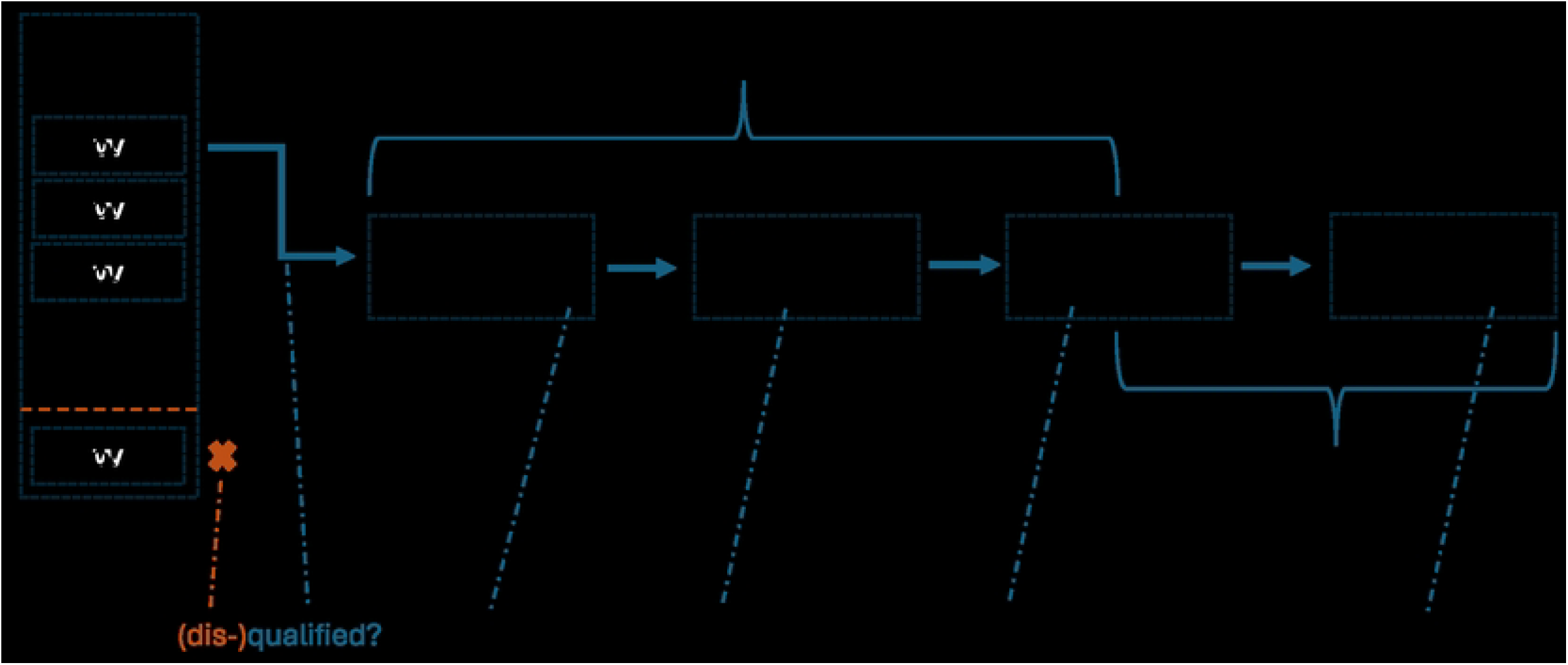
Workflow for the inspection of Boolean modelling tools. Boolean modelling tools are searched and selected. 2) Qualified tools are evaluated following usability criteria. 3) They are evaluated based on FAIR4RS criteria. 4) The tools are further evaluated on the quality and quantity of the implemented functionalities. 5) Common functionalities are benchmarked to evaluate the model size constraints.

**Fig 2.**
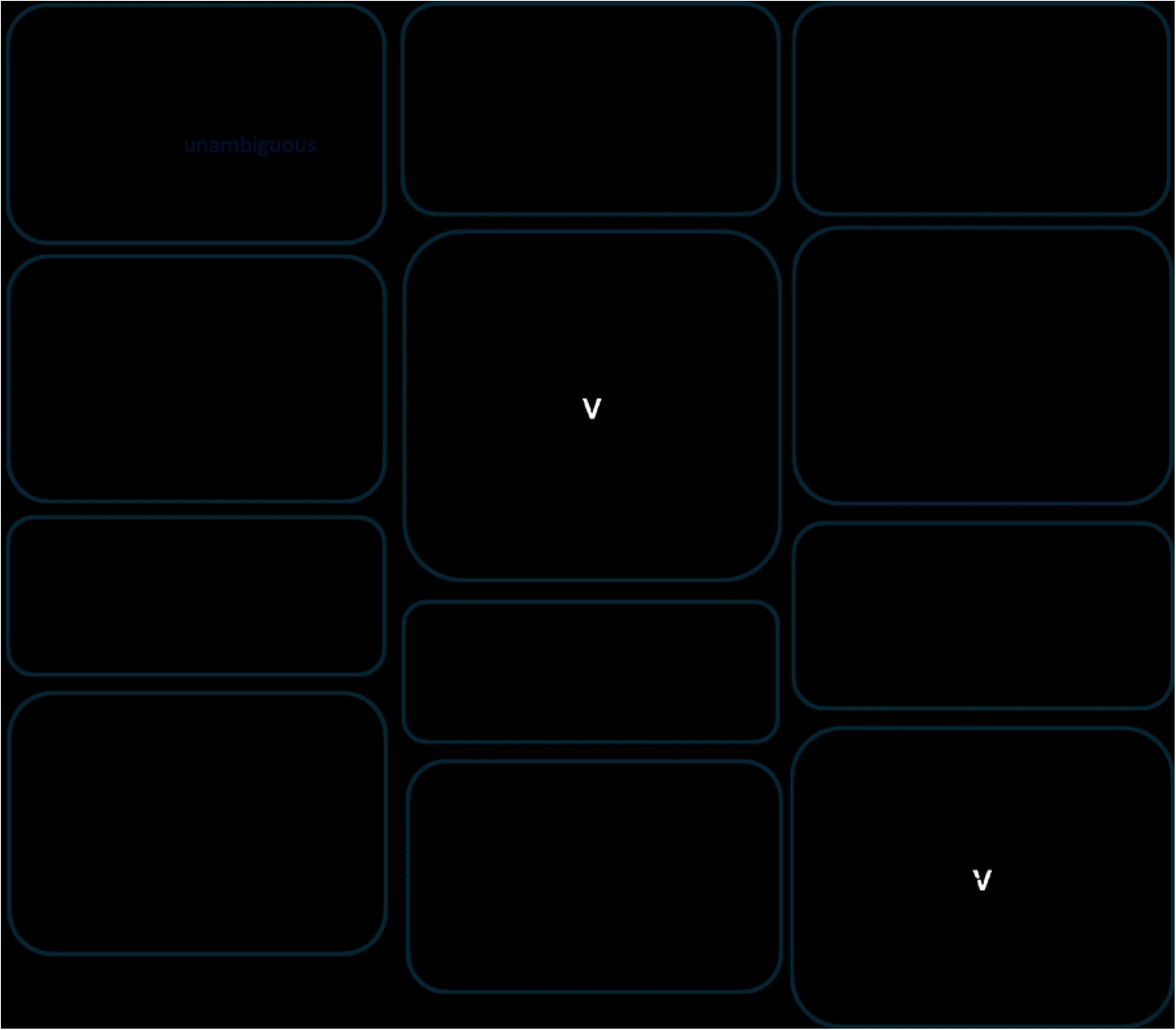
Sub-criteria for a usability evaluation. Every criterion and sub-criterion is referenced by an identifier [C.S.], in which C is the criterion identifier and S is the sub-criterion identifier.

The CoLoMoTo (Consortium for Logical Models and Tools) website, which was used as the basis for the tool selection, referenced the following tools for Boolean modelling: PyBoolNet, BoolNet, CellNetAnalyzer, Cell Collective, BioLQM, EpiLog, CellNOptR, TemporalLogicTimeSeries, LogicModelClassifier, SQUAD(&BoolSim) and GINsim [9, 11, 15–25]. The MaBoSS tool was also referenced on CoLoMoTo, for which the web version, WebMaBoSS, and the Python Interface Version PyMaBoSS were examined [26–28]. The list of tools was expanded by BooleanNet, PhoneMeS, BoolSi, GatekeepR, BooleSim and ViSiBooL, as well as the ViSiBooL Extension “Pertubation Screening v2/AutoScreenBN” [29–34]. These tools were found by literature search or referenced by the documentation of other tools. The tools were sorted into groups according to their general type of interface, and the corresponding publications along with the programming languages were collected (S1 Table).

All but five tools were runnable and accessible, and necessary information for their execution was supplied. Tools that did not fulfil these three criteria were not further investigated. Installation of both CellNOptR and PHONEMeS, when attempted following the provided documentation, proved unsuccessful, and subsequent troubleshooting efforts did not resolve the issues. The tools TemporalLogicTimeSeries and LogicModelClassifier were disqualified from further evaluation because no documentation and license information were found. Furthermore the tools had received their last update in 2014. “Squad&BoolSim” was mentioned on the CoLoMoTo website and was disqualified from further evaluation, as the service is not mentioned anymore by the Swiss Institute of Bioinformatics, and no other source for the tool’s source code was found. However, a standalone version of BoolSim was mentioned as a legacy tool by the Swiss Institute of Bioinformatics and is included in the evaluation.

### Usability evaluation

The fifteen selected tools were first classified according to their user interface type as seen in Fig 3, and then, following the evaluation criteria displayed in Fig 2, evaluated. The interfaces of tools for Boolean modelling can be broadly classified into graphical user interfaces (GUIs) and text-based forms of interaction, whereby the respective class significantly influences usability and accessibility through shared advantages and disadvantages. Tools that utilise a GUI can be delivered as a web tool, allowing access for users via a web browser of their choosing without the need for any installation of the tool itself. Alternatively, the tools can be delivered via an executable application that can be navigated by a GUI. Some downloadable tools may offer a GUI as well as a textual interface. A possible approach to the classification of textual interfaces would be to divide them into two groups, depending on whether they utilise the command line, in which case functionalities are called and parameters are defined by flags, or whether they are libraries. These extensions to programming languages supply developers and modellers with functions and objects that can be modified or integrated into larger scripts by the user. Of the 15 tools included in the usability evaluation four were classified as webtools, four as GUI tools, three command-line tools and four libraries (Fig 3).

**Fig 3.**
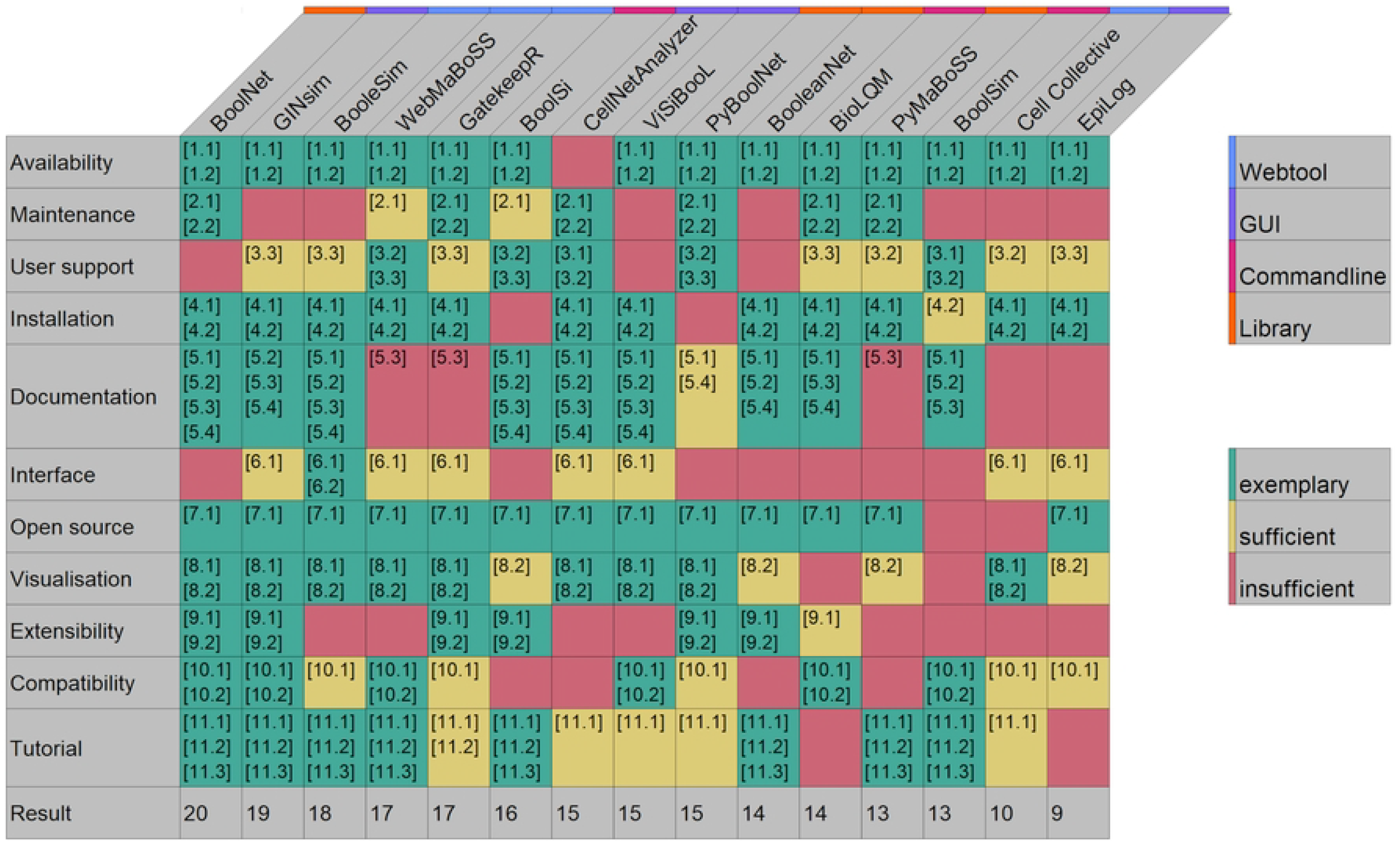
Overview of the usability evaluation for the examined tools. Fulfilled sub-criteria are noted with their accession (Fig 2). The boxes were coloured green, indicating exemplary fulfilment, yellow for sufficient and red for insufficient fulfilment. The Result row shows the number of fulfilled sub-criteria. Tools are ordered by resulting score.

As can be seen in Fig 3, the best-ranked tools were BoolNet, which fulfilled 20 sub-criteria, as well as GINsim (19) and BooleSim (18). BoolNet excelled at everything except user support and interface, since it has no GUI. GINsim’s last version being from 2018 lacking maintenance was its biggest usability flaw, while BooleSim’s lack of maintenance, in-code documentation, and modularity were its biggest usability constraints.

CellNetAnalyzer is the only tool that shows limited software availability. It was developed as a MATLAB library and requires a MATLAB license. Additionally, the software must be requested via the software’s home page, where personal information must be supplied.

The maintenance status for the tools shows a mixed landscape. Updates (inspected for the tools in March 2024) range from very recent (e.g., GatekeepR in November 2023, CellNetAnalyzer in 2023, BioLQM and BoolNet/PyBoolNet in mid-2023) to older or unknown statuses. Some tools, like Cell Collective and ViSiBooL, lack clear version or release date information, with ViSiBooL’s description only stating 2016. Other tools, including BooleSim (February 2021), WebMaBoSS (July 2022), EpiLog, GINsim, and BoolSi (all since at least 2022), show less recent activity. BooleanNet and ViSiBooL appear unmaintained since 2020 and 2016, respectively.

User support varies across the tools. WebMaBoSS offers direct email support and actively resolves GitHub issues; other web tools also address all GitHub issues, only Cell Collective has no public Git repository, but can alternatively be reached via email address. CellNetAnalyzer and BoolSim provide support via phone and email. EpiLog, GINsim, BoolSi (which also offers email support) and BioLQM have all their Git issues answered. For libraries, PyBoolNet provides support through a maintained Git or email. In contrast, BoolNet lacks user support, and BooleanNet has an unanswered Git issue from 2020, its sole contact method. Most common method for user support offered is Git Issues.

The simplicity and reliability of the installation process varies significantly across the tools. All web tools were directly accessible and required no installation. Similarly, all GUI tools were straightforward to install, typically launched via a simple code line or directly executable Java files. However, some tools presented installation challenges. BoolSi, a command-line tool, had issues with its described GitHub installation method, necessitating the use of a Docker image for evaluation. PyBoolNet also proved problematic to install, with the provided instructions failing to lead to a fully functional setup. Because of issues with the local PyBoolNet installation it was ultimately inspected within the CoLoMoTo notebook environment, where it was fully functional. In contrast, both BoolNet and BooleanNet libraries were successfully installed with minimal effort, requiring two or fewer lines of code.

Documentation quality in the sense of code external explanation of detailed information on how to install and use the given tool, varies considerably across Boolean modelling tools. BooleSim stands out among web tools with its comprehensive documentation, which includes instructions for local installation and a complete guide for usage with fitting visual aid. WebMaBoSS and GatekeepR have limited documentation, only fulfilling the criteria for guiding installation. For GUI tools, EpiLog’s documentation was unavailable, as the website was offline when inspected. The other three GUI tools generally have a complete documentation, though GINsim’s was considered incomplete and, along with its homepage, was unavailable for multiple periods during our study. In the command-line category, the tools are largely well-documented, but BioLQM’s description of possible flags and options is incomplete, and BoolSim lacks documentation for installation. Among the libraries, BoolNet’s documentation is comprehensive. BooleanNet’s GitHub README provides functional documentation, but its “Google Code” linked documentation is outdated and points to an unreachable website, with broken installation instructions. Nevertheless, the installation via GitHub was successful. PyBoolNet’s documentation is not current, and its descriptions of function parameters and inputs are incomplete.

GUIs were provided by the Web and GUI tools. BooleSim’s GUI is particularly strong, offering helpful tooltips to guide users, a unique feature among all tools. Most of the tools examined are open source, promoting transparency and community contribution. Exceptions to being open source were BoolSim, of which only binaries were available and Cell Collective, which was not available at all.

Visualisation capabilities vary greatly. All web tools can visualise networks and additional information, with GatekeepR producing a single visualisation that highlights Gatekeeper and Hub-Nodes. For GUI tools, EpiLog cannot visualise the Boolean network, while other GUI tools offer both network and further visualisation capabilities. Among command-line tools, BoolSi creates visualisations for time-series simulations, but both, BoolSi and BioLQM, do not visualise the network itself. BoolSim lacks any visualisation feature. In libraries, PyBoolNet and BoolNet can visualise both the network and the state transition graph, whereas BooleanNet only provides visualisations for time series.

Extensibility often was exemplary or sufficient, with only one tool fulfilling exactly one of two criteria. GatekeepR’s code structure is exemplary for its modularity and documentation. GINsim’s code is also very modular and well-documented. BoolSi’s code is modular and documented, facilitating modification, while BioLQM lacks a complete in-code documentation. All library tools were written modularly and are internally documented, specifically designed for use within and in combination with custom code. In contrast, ViSiBooL lacks proper code documentation and has an unclear modular structure, making code reuse time-consuming. EpiLog’s code is sparsely documented, while CellNetAnalyzer’s code lacks a clear hierarchical organisation, with many scripts residing in a single main directory alongside subfolders that do not follow a documented or consistent organisational scheme. Cell Collective is not extendable due to unavailable source code.

The variety of formats for models and the fact that none are supported by all makes it important for tools to support multiple common formats. Compatibility is a key factor in the interoperability of Boolean modelling tools. BooleSim supports four formats, including BooleanNet’s format, and BoolNet, which is the only standard format supported. WebMaBoSS is unique among web tools for supporting two standard formats (Bnet and SBML-qual), in addition to the GINsim format and its own format. For GUI tools, both ViSiBooL and GINsim demonstrated good compatibility by allowing model import in two standard formats. However, CellNetAnalyzer’s import function for SBML-qual models was non-functional during testing, it is fixed in a newer version. In the command-line category, BoolSi is limited to its proprietary data format, which restricts its compatibility with other software. Conversely, BioLQM excels in this area, supporting impressive ten formats - more than any other tool examined. These include BoolNet, BoolSim, BooleanNet, SBML-qual, MaBoSS, Multi-valued Net, Petri Net Export, CNET, GNA, GINML, and Truth Table formats, thereby significantly enhancing cross-tool compatibility by covering all standard formats and allowing the conversion of models from one to another. Among libraries, BoolNet supports both SBML-qual and its native BoolNet format. PyBoolNet handles Bnet and GINsim formats. PyMaBoSS and BooleanNet do not support any of the specified standard formats.

The provision of example models and tutorials significantly impacts user onboarding. The Cell Collective offers models by its model repository to explore all functionalities, even though no example model or tutorial is explicitly mentioned. GatekeepR provides an example model and streamlines its main use case. BooleSim and WebMaBoSS provide example data and exhaustive tutorials to help users test functionalities. Among GUI tools, GINsim is notable for its complete tutorial covering most use cases. For command-line tools, BoolSi as well as BoolSim include example data and a comprehensive tutorial, while BioLQM lacks both. In libraries, BooleanNet offers example data and a guide for its primary use case, along with tutorials for advanced features. PyBoolNet has documentation and a tutorial, but there are inconsistencies with the current version’s function and class names.

### Evaluation of FAIR4RS conformity

The FAIR for Research Software (FAIR4RS) principles adapt the established FAIR guidelines for data – Findable, Accessible, Interoperable, and Reusable – to improve the sharing and reuse of software as a primary research output [35, 36]. Nevertheless, these principles function as high-level guidelines rather than a prescriptive checklist, meaning there are no universally defined implementation rules. This inherent flexibility gives rise to ambiguity, with the result that compliance can be interpreted with varying levels of stringency. The outcomes depend on the extend to which these principles are strictly interpreted, particularly regarding the necessity for machine-readable as opposed to human-readable information.

Although human readable information was included and the FAIR4RS criteria were not assessed as strictly as possible, differences in conformity were found: PyMaBoSS, BoolNet, BioLQM and GINsim fulfilled seven to eight of the ten evaluated aspects, whereas Cell Collective only fulfilled one and a half (Fig 4). There was a lack of adequate machine-readable metadata for most tools, optimally many of the FAIR4RS criteria should be fulfilled in a stricter more machine-readable sense. Two criteria were dropped for the evaluation: FAIR4RS-Criterion A2 asks whether metadata for software will still be accessible when the tool itself is no longer available. While software repositories hosted on large public code-sharing platforms like GitHub or BitBucket are typically more permanent than privately hosted websites, these repositories can also be made private. I2 concerns the question of whether the tool has qualified references to other objects. While most tools had a reference to another object, such as a formalism used for their data type or an algorithm used, it was decided that assessing which tools sufficiently mentioned other objects and which did not was too subjective. Therefore, I2 was left out, too. If example data and documentation were available, these aspects were also analysed in the usability evaluation.

**Fig 4.**
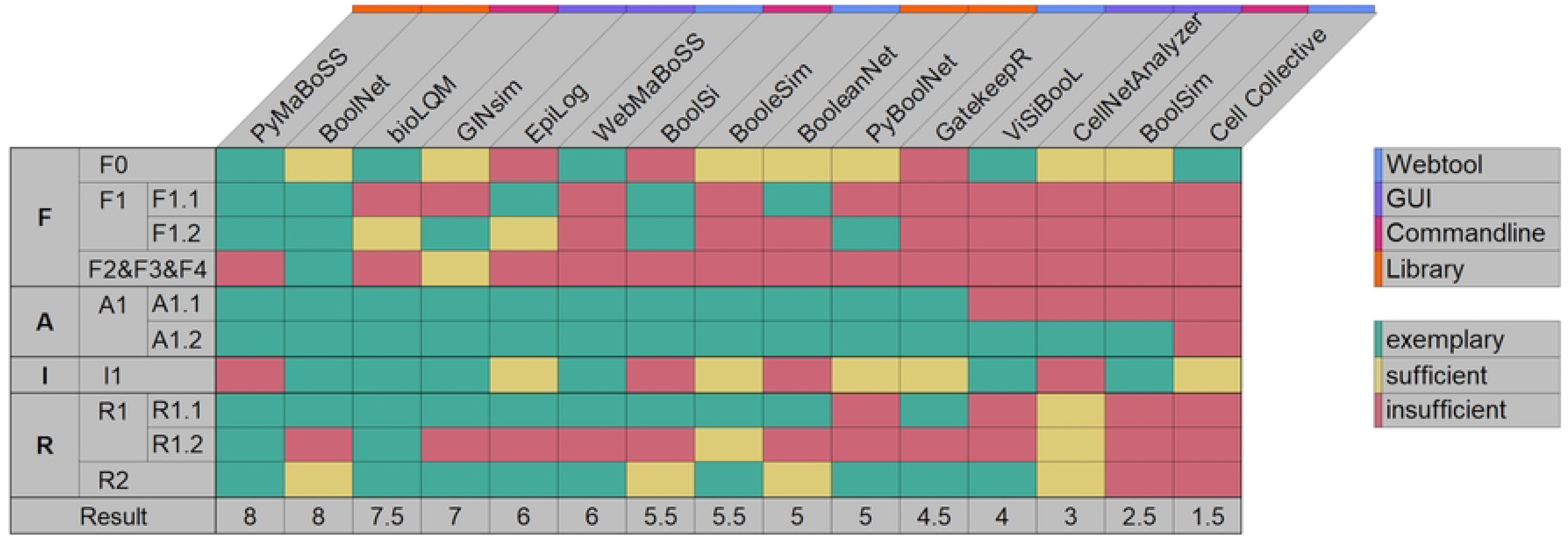
Overview of FAIR4RS conformity for the evaluated tools. The Result sums up the number of exemplary fulfilments valued as 1 and sufficient fulfilment as 0.5. S3 Table provides detailed information on the collected notes used for grading of the different items.

While FAIR focuses on machine readability, the uniqueness and persistence of a software identifier can be interpreted at the level of the software name, as well as the requirement for a DOI. Therefore, in this case we created F0 as an additional criterion. F0 refers to the test for the uniqueness and persistence of the software’s name. Most tools had a unique name, except for BoolSi, EpiLog, and GatekeepR where either a tool with the same name existed or a word was so close that they were not easily found. Multiple tools used different capitalisations which optimally should be avoided. BooleSi and BoolSim are two names that are so similar that it can lead to confusion between them, names this similar should ideally also be avoided. Principle F1.1 requires elements at different levels of granularity to have persistent identifiers. For software, this granularity can range from the entire tool, modules, functions, or even code blocks. Due to the scope of this evaluation, our analysis of F1.1 was limited to assessing the persistent identifier for the software tool as a whole. It was fulfilled, if the software had a dedicated DOI. F1.2 concerns versioning and whether the versions had a DOI and release-specific metadata. Only BooleanNet, BoolNet, BoolSi, EpiLog, and PyMaBoSS had a DOI. A few tools had proper versioning, but even fewer tools offered concrete Git releases for their versions with release notes. For some tools no version information was found at all.

F2, F3, and F4 dealt with the machine-readable metadata supplied with the tools. This metadata should be included in the software itself, but this was only the case for BoolNet. All the software was available, except for Cell Collective, which could be used as a web tool, but the software was nowhere to be found. Retrieving CellNetAnalyzer, ViSiBooL, and BoolSim required a manual download from their respective websites, as they were not accessible via a standardised, automatable procedure like a git pull command. (A1.1).

The Criterion I1 assessed if the tools could read, write and exchange data in a community standard format. This is equivalent to the Compatibility requirement from the usability criteria. Noteworthy was BioLQM, which supports 10 formats and allows to convert them into each other.

R1.1 was about having a clear, accessible and machine-interpretable licence. A licence could not be found for PyBoolNet, Cell Collective ViSiBooL and BoolSim. CellNetAnalyzer had a licence on their webpage, but it was custom and therefore not machine interpretable. R1.2 asks if the software is associated with detailed provenance. R1.2 was interpreted as meaning whether there is information about why or how the software was created, or if it was built on a predecessor. Only BioLQM, PyMaBoSS, BoolSim and WebMaBoSS provide this information. R2 concerned the question of whether relevant software was referenced enough. This was deemed sufficient if the software clearly listed its dependencies. If additionally useful software was mentioned as an extension for further analysis or data type conversion, this was considered exceptional. Tools like EpiLog and BioLQM fulfilled this criterion, as they had a pom.xml file. It would be best practise to additionally name important libraries in the documentation. Some tools also mentioned extensions and other tools that they were based on or inspired by. Cell Collective and BoolSim lacked all this information.

### Evaluation of functionality

The presence of different functionalities in the available tools was rated as complete, partial or absent (Fig 5). The categories for the functionalities of the tools were based on “The CoLoMoTo interactive notebook: accessible and reproducible computational analyses for qualitative biological networks” by Naldi et al. (2018) and expanded by the different updating schemes for Boolean models [37]. The highest number of assessed functionalities was with GINsim and Cell Collective, which both showed everything except model inference. Model inference is a specialised functionality not found with many tools. They both had synchronous and asynchronous updating but no additional alternative updating schemes. The Cell Collective lacked attractor analysis.

**Fig 5.**
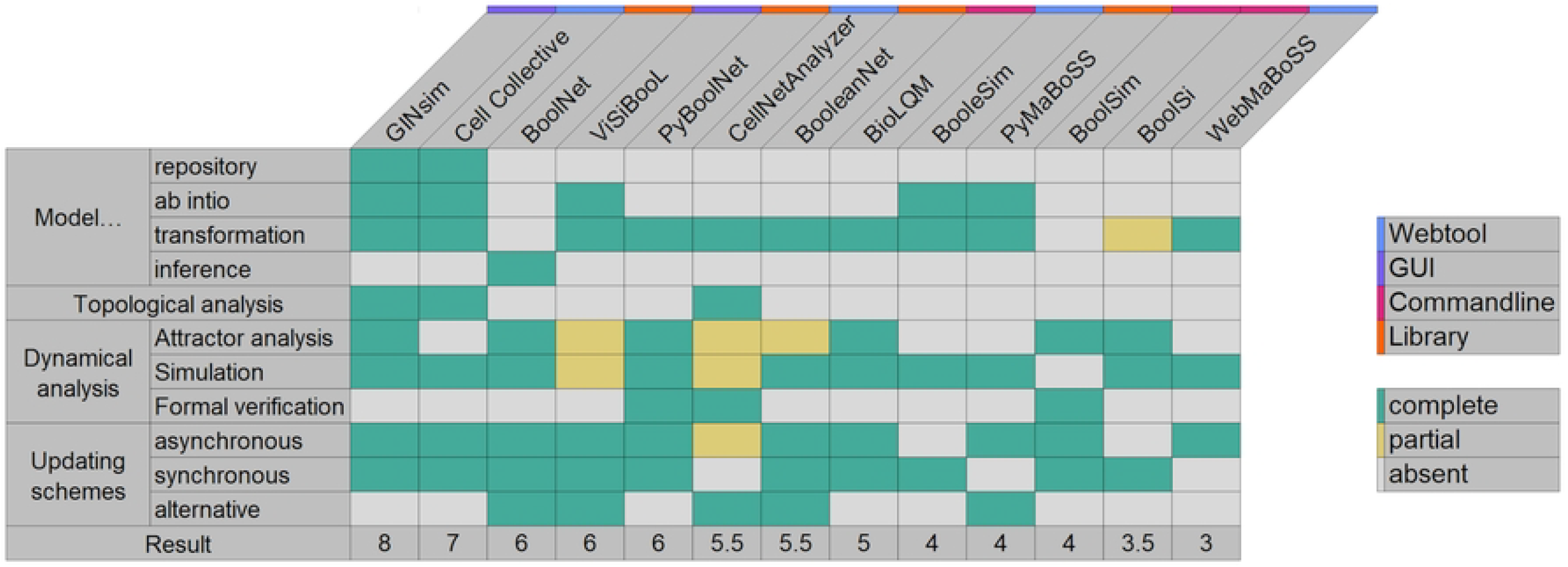
Feature matrix and characteristics of tools for Boolean modelling and analysing biological networks. “Model… “ encompasses features for creating, retrieving and modifying a model, where “ab initio” indicates the interactive construction of a model from scratch, and “transformation” involves operations like mutations, Booleanization, model reduction, etc. “Model inference” is the derivation of Boolean networks that are compatible with given properties and observational data. “Model repository” denotes searchable databases of models. “Topological analysis” involves extracting features from the regulatory graph, such as feedback cycles and graph theory measures. “Dynamical analysis” pertains to properties related to the state transition graph of Boolean or multi-valued networks. “Attractor analysis” focuses on identifying stable states, cyclic attractors, and related characteristics; “Simulation” refers to sampling trajectories within the state transition graph which may be parameterised with stochastic rates and mutations. “Formal verification and control” involves exhaustive analyses to assess strict temporal properties, such as reachability, and to deduce mutations to control the system. Figure oriented on Naldi et al. 2018 [37].

Model inference was supported by only one tool (BoolNet), while topological and formal verification were supported by three tools each. With a minimum of three functionalities WebMaBoSS focused on asynchronous simulation and statistical analyses of multiple asynchronous runs. Five tools had further updating-schemes including a transformation to an ODE model by CellNetAnalyzer and specialised rules for usage of time-delayed states by ViSiBooL. BoolNet offered alternative synchronous approaches where some nodes received updates only every few iterations.

### Benchmarking results

Boolean modelling tool’s ability to apply their functionalities is limited by the size of the model, with some tool’s functionality limited to models smaller than 25 nodes and other functionalities applicable for models larger than 10,000 nodes (Fig 6).

**Fig 6.**
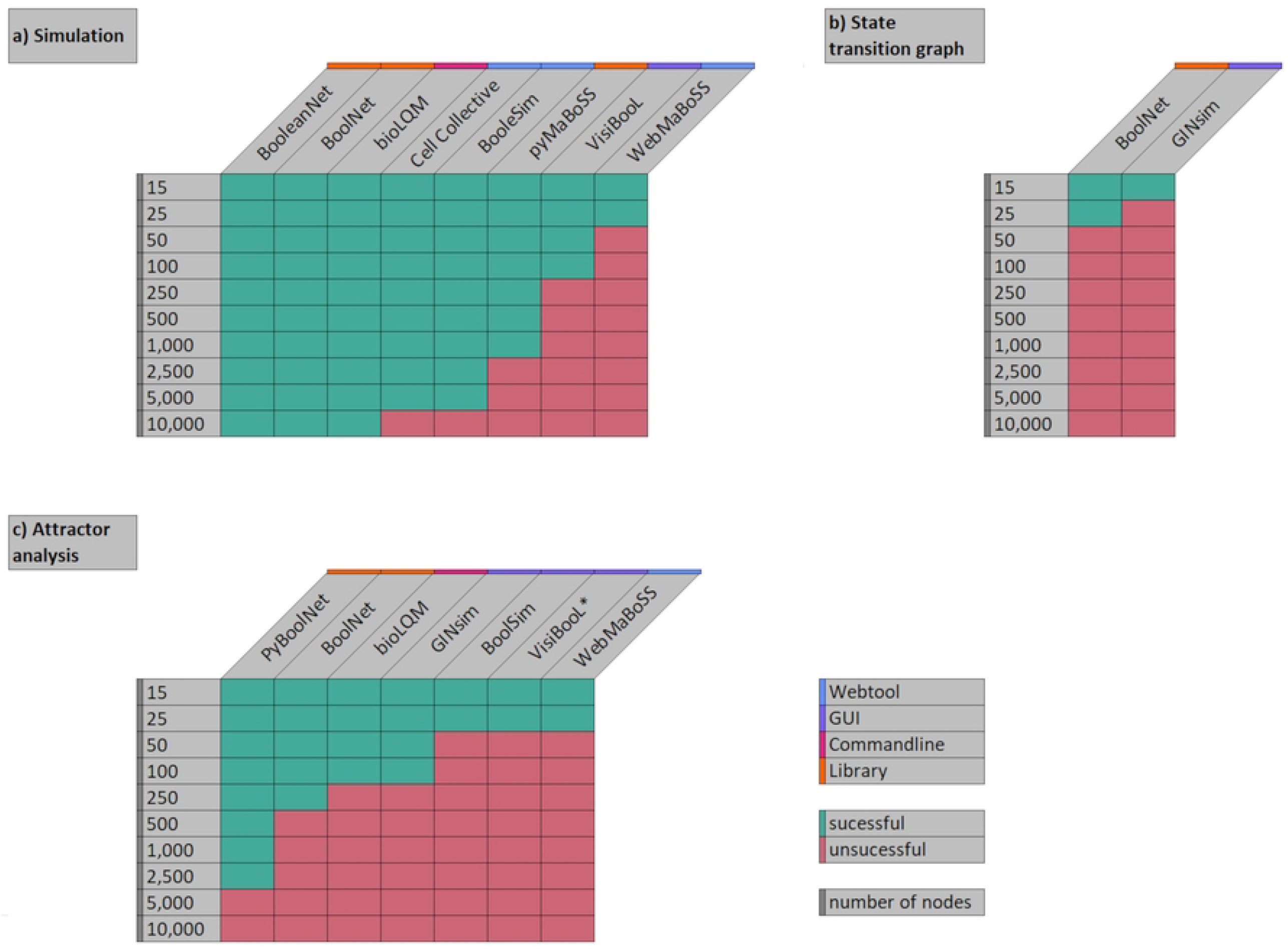
Capability of benchmarked tools to perform functionality on the models from the benchmarking set. Ability to (A) perform a simulation, (B) create a state transition graph (a comprehensive form of simulation) or (C) perform an attractor analysis for models with different numbers of nodes, while each node is controlled by two other nodes. Execution of a functionality of a tool for a given model size was deemed successful when the functionality yielded results within 20 minutes. The benchmarking included all tools for the given functionalities found in the functionality evaluation (Fig 5) for simulation and attractor analysis. BoolSi and CellNetAnalyzer are excluded, as they lack a compatible format. ViSiBooL’s attractor analysis was performed with PertubationScreening v2.

Since computational time and resources increase with model size [38], the time required to perform analytical functions becomes relevant for users modelling large systems. Therefore, benchmarking with models of different sizes was performed to assess the different capabilities of tools to apply their functionalities with large models. For conducting the benchmarking, a set of Boolean models was generated utilising BoolNet’s generateRandomNKNetwork function (S1 Code). These models ranged in size from 15 to 10,000 nodes with each node having two incoming edges from random notes. The number of incoming edges per node was set to two, as the median ratio of edges per node for the 20 newest models in the Cell Collective Repository was 1.7, which is the closest natural number.

The ability of 17 functionalities from 11 tools to simulate or analyse models of different sizes was benchmarked. The tools BoolNet, BooleanNet and BioLQM were able to simulate all the models tested (Fig 6A). BooleanNet simulated the 10,000-node model in 27 seconds, 7 seconds faster than BioLQM (34 seconds) and about six times faster than BoolNet (186 seconds) (S2 Table). BoolSi and CellNetAnalyzer were excluded, as CellNetAnalyzer’s import function was not working and BoolSi was only able to import from its own format.

The web tools BooleSim and The Cell Collective performed simulations for models up to 5,000 nodes, PyBoolNet for models up to 2,500 nodes. The Webservice BooleSim can process models up to the size of 5,000 nodes. The error occurring when loading the 10,000-sized browser models is “is lacking storage space”. The library PyMaBoSS, used as part of the ColoMoTo Notebook, was able to simulate all models smaller than and including the 1,000-node-sized models; the tool states that it is limited to models of a maximum of 1,024 nodes. The web service WebMaBoSS yielded a result for the 15 node-sized models and then 25-sized model within a few seconds but was limited to 15 output nodes. The simulation functionality of WebMaBoSS encompasses the identification of fixed points, thereby satisfying the requirement for an attractor analysis in addition to that of the simulation.

Other tools offered functionality that met the simulation requirements but calculated the entire state transition graph rather than the path from a starting state (Fig 6B). BoolNet was successful with the 25 sized model, which is in accordance with its documentation statement that 29 should be the upper limit. GINsim was only successful with the 15 sized model. In finding attractors, PyBoolNet’s function to find steady states can deal with models substantially larger than the other tools, successfully finding the attractors for the 2,500-sized model (Fig 6C). BioLQM can find attractors for models up to 100 nodes, and BoolNet can find attractors for models up to 250 nodes. GINsim was able to find stable states for models up to 100 nodes and attractors for up to 15 nodes. BoolSim, WebMaBoSS and ViSiBooL all find all attractors for models up to 25 nodes.

The largest test model, GatekeepR was found to be operational on, had 2,500 nodes, with the process requiring a duration of approximately six minutes.

### Result Summary

Our evaluation revealed that no single Boolean modelling tool provides a comprehensive solution, with every tool showing significant gaps in either functionality, usability, or adherence to FAIR4RS criteria (Fig 7). BoolNet achieved the highest usability score, fulfilling 20 out of 25 sub-criteria, while the selected tools fulfilled 15 sub-criteria on average. Each sub-criterion was fulfilled by at least one tool, on average 40% of the criteria were unfulfilled. However, the fact that even the best-performing tools left 20% of sub-criteria unfulfilled shows that there are many opportunities to improve the usability of Boolean modelling tools. Similarly, the tools account for an average of 52% of the functionalities and fulfil 53% of the FAIR4RS criteria.

**Fig 7.**
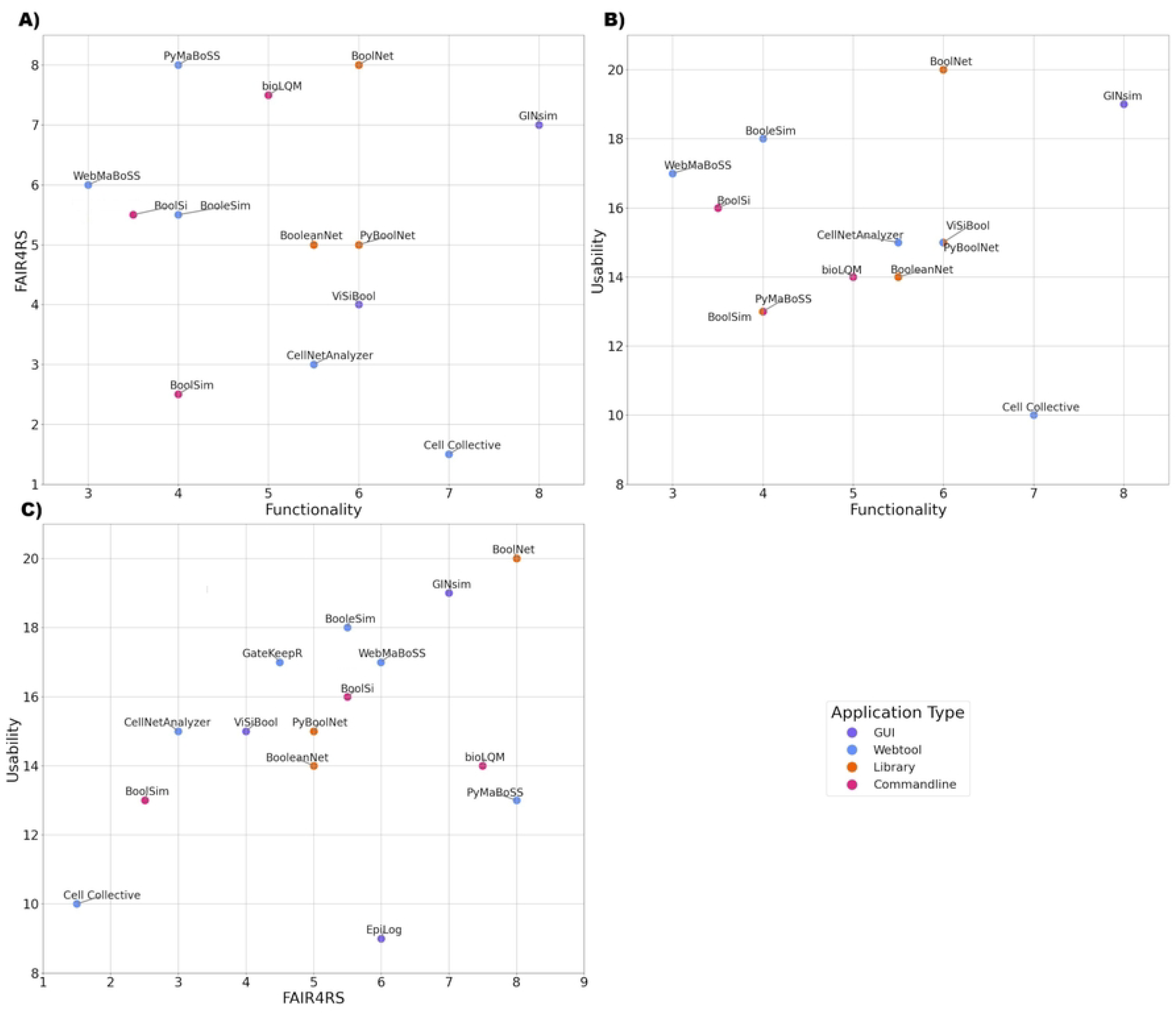
Overview of Usability, FAIR4RS-compliance and Functionality of evaluated Boolean modelling tools. For the usability axes, the number of fulfilled sub-criteria, there were 25 sub-criteria, are displayed, for FAIR4RS the number of analysed fulfilled items, there were 10, are displayed. For Functionality the number of functionalities with partial functionalities counted 0.5 with a possible maximum of 9. FAIR4RS vs Functionality (A), Usability vs Functionality (B) and Usability vs FAIR4RS (C) are plotted. All axes do not start at 0.

## Discussion

Boolean modelling tools are a diverse type of application, no tool met all the criteria from any of the four different aspects analysed. As for FAIR compliancy and usability, 40% of the applied criteria were fulfilled on average. Functions varied, as well as how the tools used common terms to classify them. Simulation capabilities ranged from tools able to simulate models with 10,000 nodes to tools that were limited to models with 50-100 or an even smaller amount of nodes.

In general, for all the evaluations that were performed it should be kept in mind that this is no exhaustive examination and that the selection of criteria as well as the benchmarking model set influences the results of the evaluation. The different criteria were not weighted, and it might be important to keep in mind that some aspects might be more impactful than others for most people. Also, even though a tool has a low score, it might still be the best choice for specialised applications.

### FAIR4RS

A strict interpretation of the FAIR4RS criteria would have been unfulfilled for almost all tools. This is mostly because the machine-readable aspect was extremely limited, metadata as a machine-readable file that came with the delivered source-code or software was only the case for BoolNet, this should be standard praxis. Older releases of a software should be available to users so that they can recreate exactly what was done by other scientists. Software should be released in stable versions and release note should be accessible to track changes. Optimally there should be a DOI dedicated for a specific release of a tool so that scientists can easily and directly cite what they used, for example by using Zenodo [39]. During the evaluation of the tools, there were instances where researcher-maintained websites were unreachable for a few days and then became available again. This issue was only observed for researcher-maintained websites, in contrast to pages on public repositories such as GitHub, which never had downtimes when examined. During the tool evaluations, the entries for GatekeepR and PertubationScreeningv2 on the ‘Uni Ulm’ website stopped being listed, making the software hard to find. Links inside the corresponding publication or website referring to the download or documentation page of a tool often led to discontinued websites. A study of papers from 2007–2017 underscores the challenge of long-term accessibility [40]. While the trend of linking to software artefacts in papers is increasing, many become unavailable over time due to broken links. In the most recent year analysed, only 58.5% of artefacts were available. With researcher-maintained websites in comparison to for example Zenondo or GitHub, websites are meant that are maintained by the academic institution themselves. When funding runs out, the institution ceases to exist or the project is taken over by other scientists, these researcher-maintained websites might be shut down or the certificates not regularly updated. Therefore it is disadvantageous to distribute tools only via researcher-maintained websites, as in these cases software and its provenance can get permanently unavailable to the research community. Public repositories on established code-sharing platforms (in combination with pulling releases to Zenodo) should be used instead, also as these often come with standardised protocols and are typically more persistently available.

### Usability and maintenance challenges

An example of where some usability criteria may not be met due to the developers’ intentions is the presence of a GUI (C.6.1). Nevertheless, having a GUI will be advantageous, particularly for users with limited programming experience who are more familiar with the biological or experimental aspects of systems biology. In addition, providing tooltips for a GUI is an important step to make tools more accessible and transparent, ensuring that users understand their analytical steps. Tooltips increase transparency and accessibility and are recommended as best practice. Only BooleSim had tooltips in its interface, a feature that the other tools lacked. Research projects provide limited staff and resources, so after funding ends, tools may not be maintained and could become unavailable. Only a fraction of the Boolean modelling tools surveyed receive regular updates. Initiatives such as de.NBI and ELIXIR provide funding to maintain software, as well as offering workshops and support, thereby prolonging the tools’ lifecycle and benefiting users. Docker is a platform that allows software to be stored in a virtual environment for users to download and run. The virtual environment includes all dependencies, even the operating system. The tool can run independently of the user’s operating system and other factors. If set up correctly, the software can be deployed and used by all users and for as long as the docker image is available [41]. Docker images were available for a few Boolean modelling tools.

For Boolean modelling tools, CoLoMoTo has created a Docker image containing 20 tools to be used within Jupyter Notebooks; this effort allows for independent testing of many tools and helps with the compatibility of the tools included in the notebook. The tools within the notebook can be used for scripting but cannot be modified. CoLoMoTo is also trying to improve code documentation for tools without documentation (“CoLoMoTo Docker GitHub Issues”). Moreover, all Git issues of CoLoMoTo are answered, and even questions about the included tools are answered. The CoLoMoTo notebook allows users to interact with the tools in a library interface type.

Documentation of code [9.2] was adequate only in 40% of analysed tools. As less than half of the tool’s code was adequately documented, peer production of a lot of these tools is severely hindered by missing documentation. Generally, more documentation also about code structure could be provided. With the emergence of more capable large language models (LLMs) in the last years, researchers have developed and trained LLMs to add documentation to code on a repository level. These and further developments can be used to add documentation to the software without much labour; in the best case, developers only need to proofread the generated documentation [42–44].

### Functionality and benchmarking

The detected functionalities for the Boolean modelling tools unveiled that most tools only offer a subset of functionalities (median number of considered functionalities: 5.5 of 11). Functionalities termed “Simulation” were implemented in different ways. Sometimes it was a time series, sometimes an attractor analysis, sometimes a stochastic overview of end states, or a state transition graph. This makes it hard for users to know tools actually deliver the functionality they need. Using the rules defined in this paper for the individual functions, an overview was created, and benchmarking was carried out for important dynamic analyses.

The ability of the tools to deal with models of different sizes varied drastically by tool and functionality for simulations ranging from fewer than 50 nodes for some tools to more than 10,000 for others. The limitations for the size of model as well as the collected computational times for completion can be seen as an orientation point, the concrete limits and computational times are subject to the system resources and the specific model provided by the users. For the tools able to do an attractor analysis with 2,500 nodes like PyBoolNet, there are multiple explanations why they are able to deal with models of larger size. They might be computationally more efficient than tools limited to much smaller size but they also might reduce the computational load by selecting a specific number of start states that are used for the discovery of attractors. Having functionalities available to analyse large models is useful, with small models of around 25 nodes, a state transition graph or an exhaustive attractor analysis may be beneficial, so that these size limited functionalities also have their niche.

### Choosing an adequate Boolean modelling tool

When choosing a modelling tool, users should first determine the size of their model and which functionalities are required. On average, 5.2 out of 11 functionalities were present in the tools examined. The 11 functionalities still exclude functionalities that are often unique to single tools. The fact that, on average, the tools showed less than half of the common functionalities highlights the importance of providing an overview of the functionalities represented in Boolean modelling tools. It is not an imperative that all tools implement all functionalities for users to have access to all functionalities. However, if tools are compatible with each other in terms of data formats and update schedules, then the utilisation of multiple tools becomes a viable approach. This reality underscores the importance of adhering to the Interoperability principle of the FAIR4RS framework. The overview we provide is designed to facilitate this process, enabling users to select a suitable tool or an effective combination of tools.

BoolNet and GINsim are the two tools we recommend for the core analytical functionalities of simulation and attractor analysis, as both tools are user-friendly and offer these functions in full, and at least for asynchronous and synchronous updates. GINsim is also one of the three tools offering topological analysis. The main limiting factor for GINsim is the model size, with a limit of 100 nodes for the attractor analysis. We therefore recommend BoolNet for the core functionality, as it was also one of the two tools with the highest usability score. It can also be used within the CoLoMoTo Notebook. BoolNet’s format is widely used with other tools, but if a specific functionality is required from a tool that does not support the format, BioLQM offers the largest number of supported formats and can be used for format transformation to make tools compatible with each other.

De novo model building with graphical editors is a problem with most tools, also BoolNet does not offer this; GINsim offers this possibility and is one of the two most user-friendly tools. An alternative is the web tool Cell Collective; these two tools are also the only ones that offer a model repository. Alternatively, manual creation inside a text editor is always possible. A user will either infer a model from, e.g., phosphoproteomics or other data or construct it manually from literature knowledge. BoolNet provides a functionality for inferring a model. Also PhoneMeS and CellNOpt are two tools for Boolean model inference. PhoneMeS extracts its information from literature knowledge and phosphoproteomics data. While PhoneMeS and CellNOpt were both rejected for evaluation as they were not functional when set up as specified by the documentation, an expert who specifically needs this functionality can invest in researching these tools and request assistance from the developers.

Formal verification and control are contained in the PyBoolNet library or CellNetAnalyzer, but with the current restrictions PyBoolNet is advised. An analysis of potential target nodes for interventions can also be done using the GatekeepR web tool. The probabilistic approach of WebMaBoSS, with its sensitivity analysis, can also lead to insight into a fitting combination of interventions. As for the functionality “minimal intervention sets”, CellNetAnalyzer should be the tool of choice for offering a complete implementation of this feature. Also the extension of ViSiBooL called “PertubationScreening v2” or “AutoScreenBN” (“Medical Systems Biology Ulm web page”), which found minimal interventions to change specific attractors but did not offer the option to specify what the new attractor should be, can be used.

BooleSim is a tool well suited for beginners who want to get their first experience with Boolean modelling; it can also be used in university lectures. BooleSim is the web tool with the highest usability score of 18. On the other hand, BooleSim scores poorly in the areas of extensibility, compatibility and maintenance, all of which are factors that do not influence a superficial use to get to know Boolean modelling; it excels in its GUI and in its documentation. Within this tool, the de novo creation of models is easy, and the state of each node is also displayed in the network view, providing an intuitive visualisation, although the creation of nodes must be done textually. The tool only supports time-series simulation and no further analysis such as attractor analysis; export of the model and generated data does not work. The tool is also limited to synchronous updating. It is an excellent choice for students experimenting with Boolean models because of its accessibility and visuality, for example as part of a lecture series.

### Criteria and their applicability to other fields

The usability criteria were partially inspired by the qualitative criteria from “A structured evaluation of genome-scale constraint-based modelling tools for microbial consortia” by Scott et al. in 2023 [14]. A core challenge in software evaluation is the transient nature of the subject. Any such assessment is inherently a snapshot in time, as tools evolve and online resources are frequently updated. This is further complicated by the difficulty of formulating purely objective criteria. Furthermore, as the evaluation in this study was conducted manually, the process is subject to human error. For example, a failure to locate a piece of information, such as a contact email, might simply mean it was not found during the search rather than it not being available.

The evaluation of code documentation is a complex undertaking. Despite the existence of standards such as Python’s PEP 8, the adherence to these guidelines remains challenging to quantify, and these standards vary across different programming languages [45]. This variability resulted in the inability to achieve a consistent and objective measurement, consequently rendering the evaluation of whether the code was ‘adequately documented’ a subjective process [9.2].

Some aspects of the established criteria are specific to Boolean models, and if the qualitative criteria are to be applied to a different set of tools, some changes to them are necessary, as, for example, the data standards have to be changed. Tool evaluators should adapt the criteria to their needs. We supply a version generalised from the version we used in this study to aid other researchers in their work, as well as a guide on how to perform benchmarking and how to apply the criteria to additional Boolean modelling tools (S1 Text). In the future the initiatives to foster good research software management should grow. The community-driven registry of bioinformatics software, bio.tools, which is led by ELIXIR Denmark, is one initiative that is advancing this [46]. Integrating FAIR4RS and usability metrics into a service like bio.tools could encourage developers to adopt good research software management practices. The main challenge would be to automate the process as much as possible.

## Conclusion

This study provides a set of developed criteria and resources for benchmarking, usability, FAIR4RS-conformity and functional comparison of Boolean modelling tools, as well as the results of their application to 20 Boolean modelling tools. It provides an overview of the Boolean modelling tools for modelling signal transduction and gene regulatory networks. It aids in finding the best tools for different use-cases including the size of the model, FAIR4RS, usability and available functions of the tools. It also illustrates the challenges and brings out issues and recommendations that should be considered in improving existing tools as well as future tools so that their overall usability and performance best suit potential users.

## Materials and methods

The supplementary material contains many notes and details on the precise use of the analysed tools. Scripts and descriptions are also provided.

### System specifications

All evaluations were carried out on a Laptop with the following specifications:

- Operating System: Microsoft Windows 11 Home
- CPU: i7-12700H, 2300 MHz, 14 Cores
- RAM: DDR5, 2*8GB, 4800 MHz
- GPU: NVIDIA RTX 3060

Webtools were accessed from this setup. Tools evaluated inside the CoLoMoTo Notebook used the CoLoMoTo Notebook Docker, which was run on this laptop.

### Tool selection

Following the evaluation workflow (Fig 1), an extensive literature review and online search was performed to uncover Boolean modelling tools applicable to systems biology (S1 Table). Main sources were the CoLoMoTo website (http://www.colomoto.org), Google and Google Scholar, as well as secondary literature and Gits.

### Filtering of tools

The tools found had to be able to perform a Boolean simulation, create a Boolean network, or have a specific analysis functionality for analysing Boolean models. The tools were also selected on the basis that they were usable, accessible and that they provided the necessary information to run them. This included the software and any necessary dependencies. It also required the information necessary to set up the tool and, if not trivial, documentation on how to use it so that the tool could be evaluated.

### FAIR4RS

Tools were evaluated using the interpretation of the FAIR4RS criteria described in the article. Aspects were checked 3 times to confirm consistency of assessment. A detailed description of the observations that were noted to decide fulfilment of the criteria can be found in S1 Table.

### Usability

Selected tools were rigorously evaluated using the usability criteria. The usability of the selected tools was analysed according to the developed criteria and sub-criteria (Fig 2). Aspects were checked 3 times to confirm consistency of assessment. The tool versions used are summarised in S1 Table. In addition to the categories from Naldi et al. (2018), categories for the available update schemes of the tools were added.

### Functionality

The categories for the functionalities of the tools were based on “The CoLoMoTo interactive notebook: accessible and reproducible computational analyses for qualitative biological networks” by Naldi et al. (2018). To increase objectivity, specific criteria were developed to define when a feature was complete, incomplete or missing.

### Benchmarking

A set of models for benchmarking was generated using BoolNet’s “generateRandomNKNetwork()” function and converted into different formats (BoolNet, Bnet, BND,BNF, SBML) using the S1 Code. For some library tools scripts to automatically benchmark were used (S2 Code). Otherwise, manual load in and functionality was carried out, measuring time from functionality start to finish, or in some cases the time for the load in (if this was the time critical step) (S2 Table).

## Supporting information

**S1 Code. Random model creation and format transformation scripts**.

**S2 Code. Benchmark scripts**.

**S1 Data. Collection of generated models in various formats for Benchmarking**.

**S1 Text. Benchmarking guide**.

**S1 Table. Overview of evaluated Boolean modelling tools**.

**S2 Table. Time data from benchmarking and tables for evaluation and benchmarking figures**.

**S3 Table. FAIR4RS evaluation overview with notes about strongpoints and shortcomings**

